# Genetic dissection of courtship song variation using the Drosophila Synthetic Population Resource

**DOI:** 10.1101/006643

**Authors:** Alison Pischedda, Veronica A. Cochrane, Wesley G. Cochrane, Thomas Turner

## Abstract

Connecting genetic variation to trait variation is a grand challenge in biology. Natural populations contain a vast reservoir of fascinating and potentially useful variation, but it is unclear if the causal alleles will generally have large enough effects for us to detect. Without knowing the effect sizes or allele frequency of typical variants, it is also unclear what methods will be most successful. Here, we use a multi-parent advanced intercross population (the Drosophila Synthetic Population Resource) to map natural variation in *Drosophila* courtship song traits. Most additive genetic variation in this population can be explained by a modest number of highly resolved QTL. Mapped QTL are universally multiallelic, suggesting that individual genes are “hotspots” of natural variation due to a small target size for major mutations and/or filtering of variation by positive or negative selection. Using quantitative complementation in randomized genetic backgrounds, we provide evidence that one causal allele is harbored in the gene *Fhos*, making this one of the few genes associated with behavioral variation in any taxon.

## Introduction

Despite a growing catalog of genotype-phenotype connections, it remains unclear what types of alleles are responsible for natural variation in most traits. We know that in some cases, such as human height, variation is explained by common alleles of small effect at a large number of loci [1,2]. In contrast, many mapped alleles in other species explain large fractions of variation or divergence in traits [3,4]. Because these latter data are ascertained with many biases, it has been suggested that mapped loci of large-effect may be the exceptions rather than the rule [5]. Supporting this hypothesis, population genetic data indicate that most adaptation is due to numerous alleles of very small effect [5]. It is possible, however, that most of this genomic response to selection has no effect on the morphology, physiology, or behavior of organisms. For example, coevolution with genomic parasites and/or compensatory evolution in response to mutation may have major impacts on the genome in ways that are important to speciation [6], but have little effect on organismal traits. The only way to resolve the genetic architecture of phenotypic variation, an important goal of both evolutionary and applied biology, is to use comprehensive, consistent, and powerful methods to connect genotype and phenotype.

In model systems, these connections have primarily been identified via quantitative trait locus (QTL) mapping in controlled crosses [3,4]. Inference from QTL studies has, however, been limited by difficulty in fine mapping QTL to identify causal genes. The *Drosophila melanogaster* community has recently tried to circumvent this problem by performing genome-wide association studies (GWAS) on ∼200 sequenced inbred lines [7]. Though this approach has successfully mapped common alleles of large effect [8], much larger samples sizes may be required for the majority of traits, where alleles may be rarer and/or have smaller effects [9–14]. In parallel to these efforts, several model organism communities have focused on developing “next generation” QTL mapping techniques that leverage technological and analytical advances to increase mapping resolution and address other challenges. To quote one recent review: “Few of the QTLs identified over the past 20 years have been resolved to individual genes, and this remains a challenging method of identifying evolved loci, although in most cases it is not clear that alternative approaches are superior” [15].

One promising approach is to start with a small but diverse panel of genotypes, mix them for multiple generations, then generate a large panel of recombinant inbred strains that allow repeated phenotyping of a single genotype: populations of this nature are now available in mice, maize, and *Arabidopsis thaliana* [16–19]. Such a population has also been developed in *D. melanogaster*: 15 strains from around the world were mixed for a remarkable 50 generations, and then ∼1700 recombinant genotypes were isolated and stabilized through an additional 25 generations of full-sibling mating [20]. Simulations suggest that this “Drosophila Synthetic Population Resource” (DSPR) has higher power and much tighter resolution than previous *D. melanogaster* mapping populations [21]. This population has recently been used to map QTL for alcohol dehydrogenase activity [20], genome-wide gene expression [22], and chemotherapy toxicity [23], but it remains to be determined how well it will perform on phenotypic traits without *a priori* candidate genes.

Male courtship behaviors in *D. melanogaster* are the focus of interdisciplinary efforts to understand the molecular basis of behavior. Impressive progress has been made on delimiting the neurons [24–27] and muscles [28–30] necessary for song production, and recent analyses have discovered variables that affect the patterning of songs relative to other behaviors [31,32]. Despite this progress, little is known about the genetic or neural control of the quantitative parameters of the song. Some of these parameters are behaviorally relevant and evolutionarily interesting, as they have diverged rapidly between closely related species, with females generally preferring the songs of conspecific over heterospecific males [33–38]. Courting *D. melanogaster* males produce a hum-like “sine song” and a more staccato “pulse song” during courtship [31,32,39]. The pulse song is likely under sexual selection, as males that are unable to produce a pulse song have greatly reduced mating success, and playing a recording of this song partially recovers this defect [33–35,40]. The pause between pulses (the inter-pulse interval or IPI; Figure 1) varies from about 30 - 45 msec in *D. melanogaster*, while IPIs of the closely related *D. simulans* are generally 45 - 70 msec [12,32,41–44]. The frequency of sound produced within each pulse (carrier frequency or CF; Figure 1) also differs between species, with *D. melanogaster* having a lower frequency pulse than *D. simulans* [46]. We have therefore focused on the IPI and CF of pulse song as evolutionarily relevant traits.

**Figure 1.**
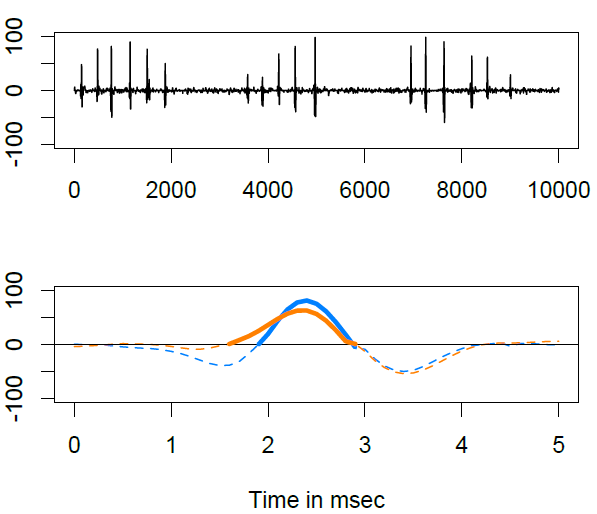
Illustrative song traits. A 10 sec interval from a single recording is shown on top. In this example, the male produced three pulse songs of 6, 5, and 6 pulses each. The distance between pulses within each song is the inter-pulse interval (IPI). A 5 msec interval from two recordings, each with a single pulse of pulse song, is shown below. The solid portion is the section used to quantify carrier frequency (CF); a 500 Hz pulse is shown in blue and a 357 Hz pulse is shown in orange.

As we discuss below, we have successfully mapped QTL explaining a large fraction of the additive variation in both IPI and CF using the DSPR. Some QTL have substantial effects, and may be useful in creating a link between genome, brain and behavior if they can be fine-mapped to causal genes and mutations. Mapped QTL for these traits are almost universally multiallelic, suggesting that the underlying genes are important regulators of these traits in nature. Using quantitative complementation in randomized genomic backgrounds, we provide evidence that variation in the gene *Fhos* underlies one of the QTL for CF, making this one of the few genes associated with behavioral variation in any taxon.

## Results

The Drosophila Synthetic Population Resource (DSPR) was started from 15 founder strains collected in Ohio, Georgia, California, and Hawaii in the United States as well as Columbia, South Africa, Spain, Greece, Israel, Malaysia, Taiwan, Peru, Bermuda, and Uzbekistan (Table S1). As shown in Figure 2, these strains differ greatly in their inter-pulse interval (IPI) and carrier frequency (CF). The DSPR was constructed by mixing strains in two sets of eight: the seven strains that contributed to population A are numbered A1 - A7, the seven strains that contributed to population B are numbered B1 - B7, and one strain, AB8, contributed to both. Over 1700 recombinant inbred strains were derived from these populations, and we measured trait values in at least 4 males from 1656 of them (N = 4-71, mean N = 16; Figure S1).

**Figure 2.**
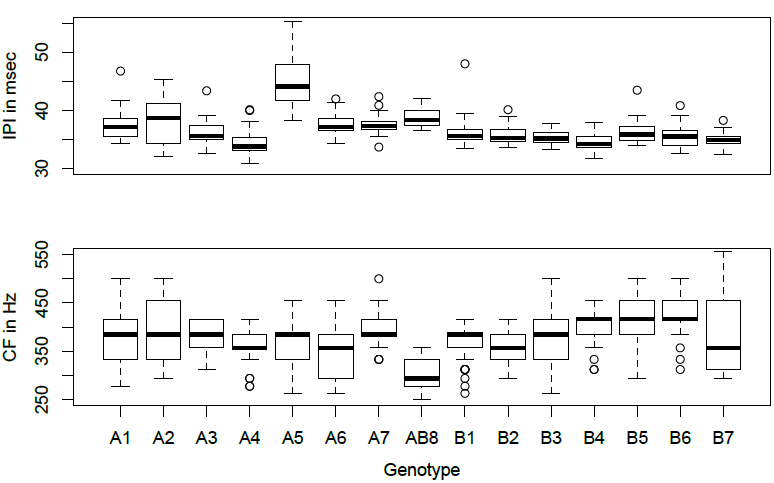
Founder phenotypes. The trait values of the 15 founder strains; box plots show the median (thick line), outer quartiles (box) range excluding outliers (whiskers), and outliers (circles). A founders and B founders were mixed separately to make the A and B populations, respectively; AB8 was included in both populations. Sample size for AB8 is only 5, as males from this founder would rarely sing in our apparatus; other lines are N = 23-46 (mean = 36).

### Heritability estimates for IPI and CF

Broad-sense heritability (*H*^*2*^), which includes both additive and non-additive genetic effects, can be estimated in the DSPR as the fraction of trait variation among recombinant inbred strains. Strain effects explained almost half the variation in IPI (46% and 45% for A and B populations, respectively) and nearly a third of the variation in CF (30% in both populations). We can also estimate the fraction of variation explained by additive genetic effects (narrow-sense heritability or *h*^*2*^) using ridge regression [47,48]. Rather than estimating the genetic variation explained by strain, this method estimates breeding values using variation among strains in the proportion of shared genomic ancestry. It is similar to methods that compare the trait correlation among relatives in a pedigree, but uses direct measurements of shared genomic ancestry rather than a historical pedigree [2,49]. We first estimated the proportion of broad-sense heritability that was due to additive genetic effects. For this purpose, we estimated trait values for each strain using one randomly chosen male, so that *h*^*2*^ is estimated on the same scale as *H*^*2*^ (see Methods) [50]. Using these estimates, *h*^*2*^ for IPI was 14% for the A population and 24% for the B population, while estimates for CF were 26% and 10%, respectively. Epistasis seems to play a variable role in these traits, as the *h*^*2*^*/H*^*2*^ ratio for IPI was 0.30 for the A population and 0.53 the B population, and for CF this ratio was 0.32 and 0.87 in A and B, respectively. For both traits, there may therefore be epistatic interactions that result from specific allelic combinations found in only one of the two populations. For the rest of our analysis, however, we focus on additive genetic effects, which explain a substantial proportion of variation and are easier to characterize.

In our QTL analysis, we can greatly increase power by repeatedly measuring males from each strain to better estimate the average trait value for a genotype. This can increase the proportion of variation explained by additive genetic effects by reducing the contribution of environmental variation to total variation. When *h*^*2*^ is estimated using the average trait value instead of only a single male per strain, the fraction of variation due to additive genetic variation increases considerably (*h*^*2*^= 31% and 40% for IPI in the A and B populations; 42% and 32% for CF in the A and B populations). Our QTL analysis also used strain means, so these *h*^*2*^ estimates are on the same scale, and thus measure the fraction of variation that can potentially be explained by additive QTL [50].

### QTL Mapping for CF

QTL mapping results are shown in Figure 3 and Table 1. For the CF trait, results were similar in the two mapping populations. At least six QTL peaks in the A population and five in the B population were apparent. To verify that these loci were all individually significant, we used forward-backward stepwise regression to reduce a model that contained a variable for each ancestral haplotype at each QTL peak. For example, at the most significant QTL in the A population, no lines we measured had ancestry from the A1, A2, or A7 founders, 443 lines had local ancestry from the A3 founder, and 46, 228, 16, and 85 lines had ancestry from the A4, A5, A6, and AB8 founders, respectively (this variable ancestor representation is due to selection and drift that occurred during the creation of the DSPR [21]). We excluded variables for ancestral haplotypes present in 10 or fewer lines resulting in a starting model with 42 variables encoding ancestry at 6 QTL in the A population. Nineteen of these variables were retained after model selection, including at least 2 significant ancestral haplotypes from each QTL (Table 1). Similarly, the final model for the B population contained 14 significant haplotypes at 5 QTL. These results indicate two things. First, all of these loci contain variation that independently associates with CF because they remain significant in a multivariate model. Second, the existence of multiple significant ancestries at each locus indicates that these QTL are multiallelic. If two variables are significant at a single locus, this implies at least 3 causal alleles, as these ancestries are significant relative to lines with all other ancestries. There may therefore be multiple alleles at a single gene underlying each QTL, multiple genes underlying each QTL, or both.

**Figure 3.**
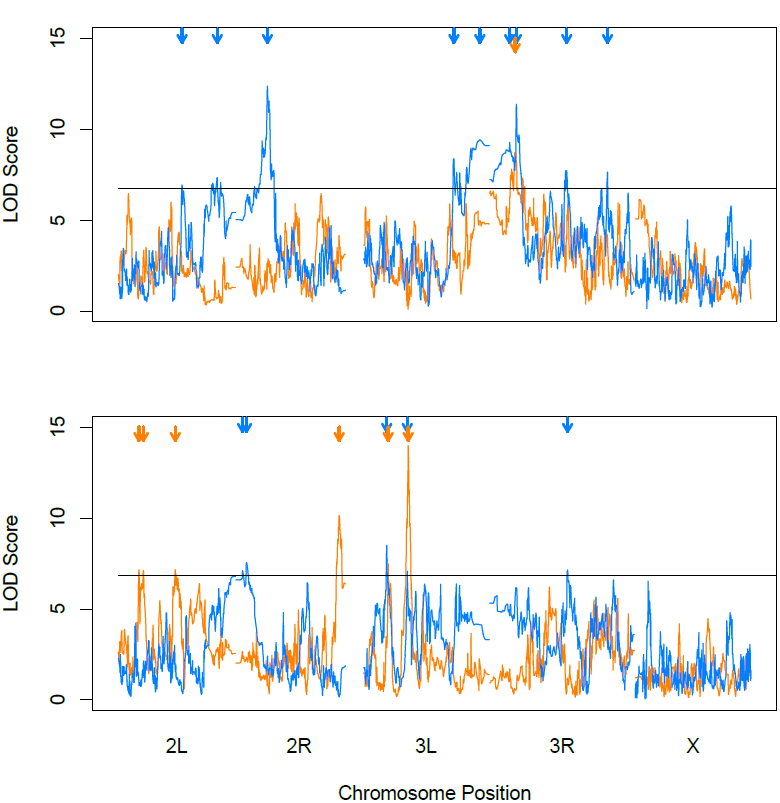
QTL mapping of IPI and CF. Genomic positions of loci affecting trait variation for IPI (above) and CF (below) courtship song parameters for the A (orange) and B (blue) populations of the DSPR. The horizontal line shows a 95% false discovery rate determined via permutation, and arrows indicate significant loci.

**Table 1.**

| Trait | Pop | QTL | % Vp | % Va | Chr | Peak (Mb) | Sig. Ancestry |
| --- | --- | --- | --- | --- | --- | --- | --- |
| IPI | A | 1 | 1.68% | 5.42% | 3R | 4.97 | A2,A3,A4 |
| IPI | B | 1 | 4.22% | 10.55% | 3R | 5.20 | B1,B4,B7 |
|  |  | 2 | 3.88% | 9.70% | 2L | 12.30 | B1,B3,B7,AB8 |
|  |  | 3 | 3.84% | 9.60% | 2R | 6.15 | B1,B3,B6,AB8 |
|  |  | 4 | 3.23% | 8.08% | 3L | 17.57 | B2,B4,B7,AB8 |
|  |  | 5 | 3.14% | 7.85% | 3R | 14.87 | B3,B4,B5,AB8 |
|  |  | 6 | 2.82% | 7.05% | 3L | 22.55 | B6 |
|  |  | 7 | 1.82% | 4.55% | 2L | 19.07 | B2,B3,AB8 |
|  |  | 8 | 1.33% | 3.33% | 3R | 3.90 | B5,B7 |
|  |  | 9 | 0.88% | 2.20% | 3R | 22.71 | B1,B4 |
| IPF | A | 1 | 10.11% | 24.07% | 3L | 8.78 | A3,A4,AB8 |
|  |  | 2 | 6.45% | 15.36% | 2R | 19.93 | A2,AB8 |
|  |  | 3 | 3.89% | 9.26% | 3L | 4.93 | A1,A4,A6,AB8 |
|  |  | 4 | 3.19% | 7.60% | 2L | 11.00 | A1,A2,A6,A7 |
|  |  | 5 | 2.98% | 7.10% | 2L | 4.87 | A3,A5,A7 |
|  |  | 6 | 2.47% | 5.88% | 2L | 4.02 | A2,A4,A5 |
| IPF | B | 1 | 7.13% | 22.28% | 3L | 4.65 | B2,B3,B5,B6,B7 |
|  |  | 2 | 3.77% | 11.78% | 3R | 15.04 | B1,AB8 |
|  |  | 3 | 2.76% | 8.63% | 3L | 8.60 | B2,AB8 |
|  |  | 4 | 2.62% | 8.19% | 2R | 1.36 | B2,B4,B7 |
|  |  | 5 | 2.06% | 6.44% | 2R | 2.13 | B4,AB8 |

These models explained a large fraction of the additive genetic variation in CF: 69% in population A and 57% in population B (Table 1). In both the A and B populations, a pair of QTL on chromosome arm 3L explained about half of this effect. The peaks of these 3L QTL in the A population are very near the peaks in the B population. This similarity in QTL location could be due to causal alleles in the AB8 founder, as this strain contributed to both populations. Indeed, AB8 ancestry is significant at both QTL in the A population and one QTL in the B population (only 5 lines had ancestry from AB8 at the other B population QTL). Additional haplotypes are significant at both QTL in both populations, however, suggesting these loci are “hotspots” of CF variation (see discussion below).

Figure 4 further illustrates the effect of one of these 3L QTL on CF (QTL 1 in the A population and QTL 3 in the B population; Table 1). When considering only this locus, median CF ranges from 356 - 399 Hz depending on ancestry, a difference of 43 Hz. For comparison, founder trait values ranged from 294 - 417 Hz, a 123 Hz range. We have previously compared outbred *D. melanogaster* (median=385 Hz, N=861) and *D. simulans* (median=500 Hz, N=936) populations with these same methods and found them to differ by an average 115 Hz.

**Figure 4.**
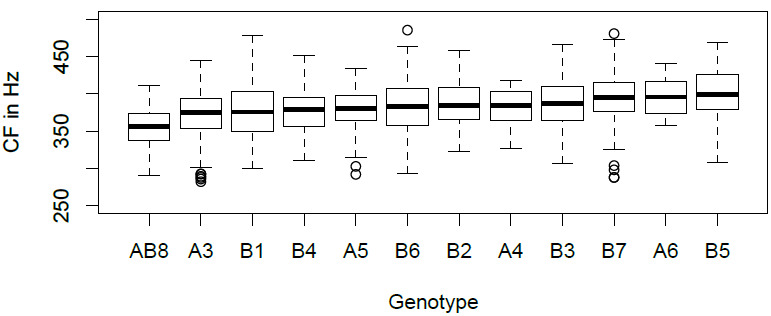
CF variation at a 3L QTL. Shown are the CF trait values for all recombinant inbred lines when grouped by their ancestry at a CF QTL located on chromosome 3L that peaks at 8.78 Mb in population A (QTL 1 in Table 1) and at 8.60 Mb in population B (QTL 3 in Table 1). Box plots show the median (thick line), outer quartiles (box) range excluding outliers (whiskers), and outliers (circles)

### QTL Mapping for IPI

Results for IPI in the B population were similar to the results for CF, in that the final model explained 63% of the additive variation (Table 1). In stark contrast, we mapped only one QTL explaining 5% of the additive IPI variation in the A population. To search for additional QTL in this latter case, we re-ran the QTL mapping program including the three significant variables at this QTL as covariates, but this produced no additional significant QTL at our 5% FDR threshold. Part of this difference between the A (one QTL) and B (nine QTL) populations appears to be due to the stochastic loss of alleles from the AB8 founder. Five of the IPI QTL in the B population had significant effects of AB8 ancestry (Table 1), which is also expected to be associated with IPI in the A population. Further investigation reveals that few of the lines we measured in the A population have AB8 ancestry at these QTL: only 30, 13, 7, 0 and 11 A lines have AB8 ancestry for QTL 2, 3, 4, 5, and 7 respectively from the B population, as numbered in Table 1. This is consistent with previous reports that founder AB8 is poorly represented in the A population, and means that little power is available to map these rare alleles in this population [20]. It seems that the genetic architecture of IPI differs between the A and B populations in additional ways. Nine QTL are significant in the A population, and only five of these involve AB8 ancestry. At all five that do, additional ancestries are also significant. The fact that our mapping results for CF were similar in both populations suggests that this difference for IPI is specific to the trait rather than the populations in general. We can also exclude epistasis as an explanation, as we are specifically estimating the proportion of additive variation explained by these QTL. This therefore seems to reflect a difference in effect sizes of causal alleles among A and B founder strains.

Though only one IPI QTL is significant in the A population, it overlaps the most significant IPI QTL (QTL 1) in the B population (Table 1). This is not due to alleles from the AB8 founder, as only one line in the A population has AB8 ancestry at this locus, and AB8 was not significant in the B population. Variables representing ancestry from Columbia, Spain, and South Africa are significant at this QTL in the A population, while haplotypes from Bermuda, Malaysia, and Israel are significant in the B population. Like the overlapping QTL for CF, it seems that this locus is a “hotspot” for IPI variation due to multiple alleles and/or multiple genes in close proximity. Simulations show that a “2-LOD drop interval” around a QTL peak is an estimate of the 95% confidence interval for the location of a causal gene, though the probable violations of model assumptions means that this is an estimate only [51–53]. The 2-LOD interval at this QTL is 660 kb in the A population and 410 kb in the B population. The overlap of these intervals spans 170 kb.

In the well-annotated *D. melanogaster* genome (version 5.56), this interval contains only 21 protein-coding genes and one lincRNA. Figure 5 displays the IPI values for all recombinant lines, grouped by ancestry at this shared 3R QTL. Median IPI ranges from 33.8 - 36.3 msec based on the genotype at this one locus: a 2.5 msec spread. Median IPIs for the founder strains varied from 33.8 - 42.0 msec, an 8.2 msec range. For comparison, our outbred *D. melanogaster* and *D. simulans* populations differ by 16.5 msec (*D. melanogaster =*35.0 msec, N=861; *D. simulans*= 51.5 msec, N=936).

**Figure 5.**
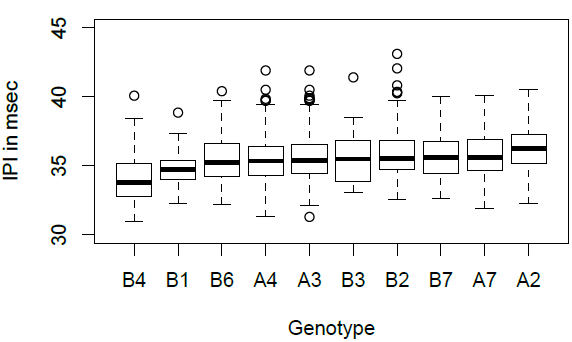
IPI variation at a 3R QTL. Shown are the IPI trait values when all recombinant inbred lines when grouped by their ancestry at an IPI QTL located on chromosome 3R with a peak at 4.97 Mb in the A population and at 5.20 Mb in the B population (QTL 1 for both populations in Table 1). Box plots show the median (thick line), outer quartiles (box) range excluding outliers (whiskers), and outliers (circles)

### Gene-level validation

Published simulations suggest that QTL effects may be due to the sum of many small effect alleles in linkage disequilibrium [5,54]. This seems less likely in our case, as the 75 generations of DSPR construction, >1600 derived lines, and extremely dense genotyping have resulted in QTL with much tighter resolution than most previous studies. None the less, some (but not all) of the mapped QTL lie near centromeric regions where linkage disequilibrium is especially likely [55]. To investigate this issue further, we explored variation in our QTL of largest effect at 8.78 Mb on 3L (CF QTL 1 in the A population; Table 1). This QTL explains 24% of the additive genetic variation in CF in population A, and is overlapped by a population B QTL (QTL 3; Table 1) at 8.60 Mb. This QTL is not near a region of low recombination [56].

The 2-LOD drop interval at this QTL is only 185 kb in the A population. One lincRNA and 25 protein coding genes are within this region (Figure S2), and many of these genes seem unlikely to be involved in courtship song. Four are structural constituents of the egg chorion, four (including the lincRNA) are expressed nearly exclusively in testes, seven are expressed nearly exclusively in malpighian tubules, and one is an enzyme inhibitor found only in the gut (as annotated at flybase.org). Four of the remaining ten genes lack any annotation and have poorly characterized gene expression profiles. Of the remaining six genes, three seem the most likely candidates: *Paramyosin (Prm)*, one of the primary structural constituents of invertebrate muscle [57,58], *Fhos*, recently implicated in muscle cell homeostasis [59], and *division abnormally delayed (dally)*, a heparan sulfate proteoglycan binding protein in signaling pathways with highly pleiotropic functions [60,61]. The 10-kb window with the maximum LOD contains most of the exons of *Fhos* and nothing else except the enzyme inhibitor expressed in the gut.

The overlapping QTL in the B population has a 2-LOD drop interval of 320 kb, but only overlaps the A population QTL for a 40 kb span. This overlapping interval contains only *Fhos, Prm*, three unannotated genes, chorion genes and two of the testes-specific genes. The small number of genes in these high-probability intervals makes it likely that variation in a single gene could have large effects on CF variation. We consider *Prm* and *Fhos* to be the most likely candidates due to their known effects on musculature.

Courtship song is generated when the flight muscles extend the wing and flex the thorax, causing the wing to “twang” [28,29]. The quantitative parameters of courtship song may be affected by variation in specific muscles: silencing motor neurons extending to the *ps1* muscle changed CF and pulse amplitude without affecting other song parameters, while other muscles had different and specific effects [30].

The *Fhos* gene at the peak of this QTL is 45 kb long (mostly introns) and has 9 annotated splice forms. Validating the role of such a complex gene is an intimidating prospect, but the quantitative complementation test provides a possible route [62].

Genetic complementation is used in molecular genetics to determine if recessive mutations with the same phenotype are alleles of the same gene [63]. The quantitative complementation test is designed for use in natural strains with different alleles at multiple loci affecting the trait of interest [64,65]. To use this test, natural strains with putatively different alleles at a gene are crossed to a loss-of-function mutation in that gene and a control strain. In the loss-of-function F1s, the effects of natural alleles at that locus will not be masked by any other allele; in the control cross, the natural alleles will combine with the control allele. If there is a significant statistical interaction between the loss-of-function mutation and the natural strains, this supports a hypothesis that natural variation at that locus affects the trait of interest. As traditionally implemented, this test can suffer from a high false positive rate due to epistasis. Natural genotypes are different at many loci, and any of these differences could interact epistatically with the loss-of-function allele to produce a false positive [66]. This problem can be greatly alleviated using recombinant inbred lines because any given allele is present in many different genomic backgrounds [67,68]. A significant interaction term therefore constitutes a high standard of evidence in cases where the genomic background is controlled or randomized. False negatives are still a problem: if the control allele combines additively with the natural alleles, for example, a significant interaction is not expected.

To investigate the role of *Fhos* alleles in CF variation, we crossed 44 DSPR strains to a strain heterozygous for a lethal *Fhos* mutant. These strains have different combinations of founder genotypes across the genome, but 15, 14, and 15 of them have ancestry from the A5, A6, and AB8 founders, respectively, at this particular QTL. Strains with A6 ancestry at this QTL have high CF, strains with A5 have intermediate CF and strains with AB8 have low CF (see Figure 4). If there are functional differences between *Fhos* alleles in these strains, we expect to see a significant interaction between the QTL genotype and the *Fhos* mutant in the F1 strains. As shown in Figure 6 and Table S2a, we found just such an interaction (p= 0.0386). Further investigation of this interaction revealed no significant differences between A6, A5 and AB8 strains with the control allele (p= 0.98; Table S2b), but highly significant differences with the loss-of-function allele (p=0.002; Table S2c). When paired with the loss-of-function allele, strains with A6 ancestry at *Fhos* had significantly higher frequencies than those with A5 and AB8 ancestry, as shown in Figure 6 (A6 vs. A5: Least squares mean difference= 4.74 Hz, p= 0.0019; A6 vs. AB8: Least squares mean difference= 4.61 Hz, p=0.0028; both p-values significant after sequential Bonferroni adjustment). This is consistent with the QTL analysis, which found that A6 ancestry at the QTL was associated with higher CF (Figure 4). The QTL analysis also found that strains with A5 ancestry have CFs intermediate to those with A6 and AB8 ancestry, but our quantitative complementation revealed no differences in CF resulting from a single copy of the A5 and AB8 alleles at *Fhos*. This may indicate that differences between A5 and AB8 alleles are strictly additive, or that they are caused by a different gene in the QTL.

**Figure 6.**
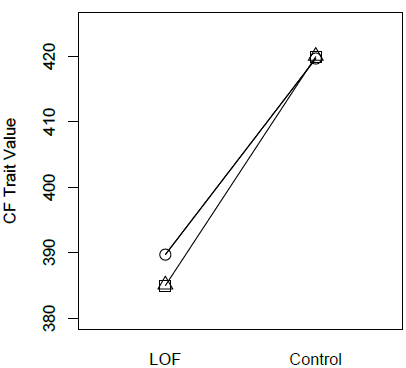
Quantitative complementation test of *Fhos* for CF. Mean CF trait values for strains with A5 (squares), A6 (circles) or AB8 ancestry (triangles) at *Fhos* combined with either a loss-of-function (LOF) mutation in *Fhos* or a control allele. The significant interaction between ancestry and the loss-of-function/control alleles indicates that ancestry does not affect CF when paired with a control allele, but has a significant effect on CF when paired with the loss-of function allele, with the A6 allele producing higher CFs than the A5 or AB8 alleles (see Table S2). N= 69-130 (mean N = 105). Plotted are the least squares means from an ANOVA performed using untransformed measures of CF.

## Discussion

Using one of the largest QTL mapping populations yet constructed, we mapped a moderate number of QTL affecting courtship song variation. Together, these QTL explain a large fraction of the additive genetic variation in inter-pulse interval (IPI) in one population and carrier frequency (CF) in both populations. QTL effects are modest compared to most well-characterized case studies [4], but large enough that we may hope to map them to specific genes and perhaps mutations. We must be cautious making interpretations about genetic complexity from QTL mapping alone, as a single QTL could be due to the combined effects of many genes in linkage, and QTL effect sizes are likely inflated [5,69]. The former issue seems unlikely for some of our largest-effect QTL due to the small number of plausible genes at these loci. Our data do support a role for multiple causal mutations, however. The QTL we mapped were not significant because of an allele from a single founder strain, but were instead due to alleles inherited from several founders. Most QTL mapping is performed with crosses between only two strains and cannot detect such effects, but recent studies in outbred populations and multi-parent RILs have also found that multiallelism is common [22,70]. This is perhaps not surprising: if the expression level of a gene affects the trait, there may be a series of alleles with variable levels of expression. This observation is very interesting, however, when considered in light of the “hotspot hypothesis” [4,15]. This hypothesis starts with the observation that repeated cases of trait evolution between species have been found to involve the same genes. This could be due to a small mutational target size: if only a few genes can alter the trait in question, these genes would be repeatedly used. Though non-random mutation is undoubtedly part of the story, the differences in gene reuse in natural populations vs. human induced mutations may also indicate a major role for natural selection [15]. In this scenario, mutations at most loci affecting a trait have deleterious side effects, so that mutations at only a few loci can pass through a selective filter and cause trait evolution.

In our data, we see that a large fraction of trait variation in IPI and CF maps to a few small regions of the genome, but that these regions almost always contain multiple causal alleles. Only one QTL is significant for IPI in the A population, but it overlaps the most significant QTL in the B population. The most significant CF QTL in the A population overlaps the third most significant CF QTL in the B population, and the most significant CF QTL in the B population overlaps the third most significant QTL in the A population. Within each population, the effect of each QTL is also the sum total of multiple alleles. It seems likely that selection plays a role in this pattern. If these alleles are nearly neutral, it is possible that there are only a few genes in which mutations that affect these traits are tolerated by selection. If these alleles are under positive or balancing selection, mutations in other genes that might affect these traits may be constrained by pleiotropy. In either case, this pattern may indicate that variation in complex traits, like divergence in simpler traits, is more predictable than previously recognized.

Finally, our results suggest that QTL mapping will play a major role in future efforts to connect genotype and phenotype, despite the current popularity of genome-wide association studies (GWAS). Our previous efforts to identify the genes responsible for IPI variation using GWAS in ∼160 inbred strains were largely unsuccessful [12].

Combining these data with data from an evolve and resequence study resulted in some progress [12,44], but considerably less than we have made here using the DSPR. QTL mapping has several major downsides, however, including 1) the difficulty in fine mapping QTL to genes, and 2) the unclear relationship between variation in the mapping population and variation in nature. In the first case, we suggest that advances in genomic manipulation make this problem tractable. We have used an existing mutation in the gene *Fhos* to identify this gene as the first known to affect CF, and one of the few known to affect natural behavioral variation in any taxa. This approach has limitations, however, as negative results are uninformative, and it is unclear how to use this test to estimate the proportion of variation explained by a gene. We are currently following up on these results using induced variation in other ways that may allow us to estimate the effect sizes of individual genes.

Our results also illustrate the second challenge of QTL mapping: making inferences about natural populations from mapping populations. We mapped IPI variation in two very similar populations and obtained very different results. In one case, we mapped 9 significant QTL that together explain 62% of the additive genetic variation in this reduced-complexity population. In the other population we could locate only one QTL explaining 5% of the additive genetic variation. Although we made no attempt to map epistatic effects, our comparisons of *H*^*2*^ and *h*^*2*^ found that these effects differ among our populations as well. The inconsistencies between our two mapping populations demonstrate the potential limitations of QTL mapping: what can we learn about the variation underlying multiple traits in natural populations if we can't even extrapolate between our A and B populations for a single trait? In this respect, we find our “hotspots” of courtship song variation very encouraging, despite the fact that these traits likely involve many other loci (both in nature and in these populations) and will be complicated by environmental effects and gene-by-environment interactions. If a small number of genes are repeatedly responsible for variation in these traits in the DSPR, it is likely that these genes (or homologous ones) will play major roles in variation, divergence and adaptation in nature. Understanding why they these genes are central to these traits could lead to insights regarding the maintenance of genetic variation and the nature of the gene-brain-behavior map.

## Methods

### D, melanogaster strains and maintenance

The founder strains used to start the DSPR were obtained from Stuart Macdonald (University of Kansas). Strains were collected from diverse locations, mostly in the 1950s and 1960s, and have been reared in laboratory conditions since that time (Table S1).

Recombinant inbred lines (RILs) were obtained from Anthony Long (University of California Irvine). Females from the wild type line RAL-380 (Bloomington stock 25189) were used as standardized courtship objects. We used Bloomington stock 11540 as an *Fhos* mutant strain, as described below. Reported values from outbred *D. melanogaster* are from a population made by mixing the RAL inbred strains collected in North Carolina [71], as described previously [45]. Trait values reported for outbred *D. simulans* are from a population founded from 500 females collected from Ojai, CA by the authors, and recorded after a single generation of lab culture. All fly strains were maintained in 25x95 mm vials on cornmeal-molasses-yeast medium in standard Drosophila incubators at 25°C under a 12-h light/dark cycle.

### Courtship song recording and measurement

We recorded courtship songs from males of the 15 founder lines and 1656 DSPR RILs when paired individually with RAL-380 females. We collected males for recording in groups of 10 under light CO_2_ anesthesia and held them at 25°C for 3-5 days to recover before recording. We collected female courtship objects as virgins in groups of 20 using light CO_2_ anesthesia and used them for recording the following day, as 1-day old females are courted vigorously but rarely copulate in our 5-minute recording interval.

Song recording hardware was adapted from an apparatus built by the Dickson lab [26], and has been described in detail previously [44]. Each male was recorded for 5 minutes, which resulted in an average of ∼200 song pulses per individual (recordings with fewer than 20 pulses were discarded). Inter-pulse intervals (IPIs) between 15 and 100 msec were considered pauses within a song bout, rather than between song bouts; the median of these values was used as the IPI for that individual. The average IPI of all RILs, 35.8 msec, is in agreement with reported values for *D. melanogaster* from other laboratories [32]. The carrier frequency (CF) has previously been calculated using either the Fourier Transform [46,72] or by measuring the zero-crossing rate [72–74]. Pulses last only a few msec (Figure 1), and we found that Fourier Transform results were inconsistent given the level of background noise in our recordings. We therefore estimated CF using the zero-crossing rate. We focused on only the highest amplitude half-cycle (Figure 1), as this was the least affected by background noise and resulted in the most consistent measurements for each genotype. To measure CF, we doubled the half-cycle time to estimate the duration of each cycle, and determined the number of these cycles per minute (hertz). As for IPI, the median value of an entire recording was used as a male's trait value. Though this method of estimating CF yielded consistent results for each genotype (see heritability estimates in Results), the averages were higher than values estimated in other labs [32]. Trait values are comparable to those previously measured in our lab using the same methods.

### Linking genotype and phenotype

Slight but significant deviations from normality were found for both traits, so data were t-rank normalized using the t.rank() function in the R package multic [75]. As shown in Figures S3 and S4, this had very slight effects on trait distributions.

Broad-sense heritability was estimated as the variance explained by strain using the lm() function in R. Narrow-sense heritability was estimated with ridge regression using the rrBLUP R library [47,48]. We first compared estimates of narrow and broad-sense heritability. Because our estimate of broad-sense heritability includes variability among flies within a strain in the total variance, narrow-sense heritability was first calculated by running rrBLUP on trait values estimated by randomly selecting a single individual from each strain (following [50]). We also estimated narrow-sense heritability on the same scale as our QTL analysis by re-running rrBLUP using the average trait values for each line, as these were the values used in QTL mapping. This latter measure estimates narrow-sense heritability as the fraction of variation explained by genotype after error variance is decreased through repeated measurements of traits. In both cases, we included subpopulation as a covariate to account for the fact that the RILs from each population (A or B) were created from two separate mixing cages (subpopulations A1 and A2 or B1 and B2), and allele frequencies may have diverged slightly among subpopulations. This is analogous to population structure in natural populations, but is easy to account for because the history of these lines is known. For each estimate, we ran rrBLUP 500 times, each time sampling 40% of the population as a training set and estimating variance explained using the other 60%; reported narrow-sense heritabilities are the means of these 500 estimates.

We mapped QTL with the DSPRqtl R package: this software is based on R/qtl [76], but was designed specifically for the DSPR population. As described in detail elsewhere [20], this package performs a multiple regression of trait value on ancestry probabilities (as estimated with a Hidden Markov Model) in 10 kb windows across the genome. The resulting *F*-statistic is then converted into a LOD score, and significance is estimated using permutation. QTL with LOD scores greater than the most significant value in 95% of permutations were considered significant, providing a 5% false discovery rate (FDR).

To estimate the combined effects of mapped QTL, we conservatively discarded some QTL found by DSPRqtl because of their close proximity to a more significant peak: loci included are shown in Figure 3 and Table 1. For each population (A or B), we started with a model that included a variable for subpopulation and one for every founder at each locus. At many loci, however, alleles from some founders were rare or absent due to drift or selection that occurred during the creation of the DSPR. We discarded variables for all founder ancestries found in 10 or fewer lines. We then fit a model using the lm() function, and reduced it with forward-backwards stepwise regression. This was done using the stepAIC() function in the MASS R library [77]. To estimate the effects of each retained variable, we used the drop1() function, which provides type III marginal sum of squares rather than the R default type I sequential sum of squares.

### Gene-level validation

To validate the role of *Fhos* in CF, we performed quantitative complementation using strain 11540 from the Bloomington Drosophila Stock Center. This strain has the genotype P{PZ}Fhos^01629^ ry^506^/TM3, ry^RK^ Sb^1^ Ser^1^, where the P{PZ} is an transposable element insert generated by the Berkeley Drosophila Genome Project [78]. As annotated at flybase.org, this insertion is located between base pairs one and two of the first untranslated exon common to 6 of 9 *Fhos* transcripts. This insertion is likely a loss-of-function, or at least a hypomorph, as it is homozygous lethal and previous investigation found that *Fhos* transcripts were reduced to barely detectable levels in this mutant [79].

To perform quantitative complementation, strains with different natural alleles at a gene of interest are crossed to a control strain and a strain containing a loss-of-function mutation at that gene. Loss-of-function F1s allow the natural alleles to be functionally hemizygous, while in control F1s the natural alleles will be expressed with control alleles. A significant statistical interaction for trait values between the F1 genotype (control or loss-of-function) provides support that this gene influences the trait under study. We crossed the *Fhos* strain to 44 DSPR strains. The mutant strain is heterozygous for a lethal *Fhos* mutation that is held over a third chromosome balancer, so the *Fhos* allele on the balancer served as our control allele. Of the 44 DSPR strains we used, 15, 14, and 15 of them have ancestry from the A5, A6, and AB8 founders, respectively, at this particular QTL, with random combinations of founder genotypes across the remainder of the genome. We recorded courtship songs over multiple days (experimental blocks) for loss-of-function and control F1 males from each of these 44 lines (N= 69-130, mean N= 105). Treatment of experimental flies and courtship song processing was identical to that described above for founder males and DSPR line males.

To test for a significant interaction between F1 genotype and founder ancestry at this QTL, we performed a multifactor ANOVA with experimental block, F1 genotype (loss-of-function or control), ancestry (A5, A6 or AB8) and their interactions as main effects. We also included the interaction between strain and F1 genotype nested within ancestry to account for any epistasis with the loss-of-function mutation among strains within a given ancestry. Interactions that were highly insignificant (all p>0.50) were removed from the model. To further investigate our focal, significant interaction, we performed the same analysis separately for each F1 genotype (control and loss-of-function). Although there were slight but significant deviations from normality in this data set, ANOVA is generally robust to minor deviations at such large sample sizes.

Nonetheless, we performed all tests using both raw and t-rank normalized data (as described above) to ensure our conclusions were valid. The test results were nearly identical (Table S2), so values reported throughout the Results are from the analysis using raw data.

## Acknowledgements

The authors thank many UCSB undergraduates for their assistance, Elizabeth King for analysis tips, and Tony Long and Stuart Macdonald for contructing the DSPR and sharing it with us. This work would not have been possible without the community supported resources available at Flybase.org and the Bloomington Drosophila Stock Center. Funding was provided by the University of California Santa Barbara, the National Institute of Health (R01 GM098614), and the National Science Foundation's support to the Center for Scientific Computing at UCSB (NSF MRSEC DMR-1121053 and NSF CNS-0960316).

---

**Figure S1.**
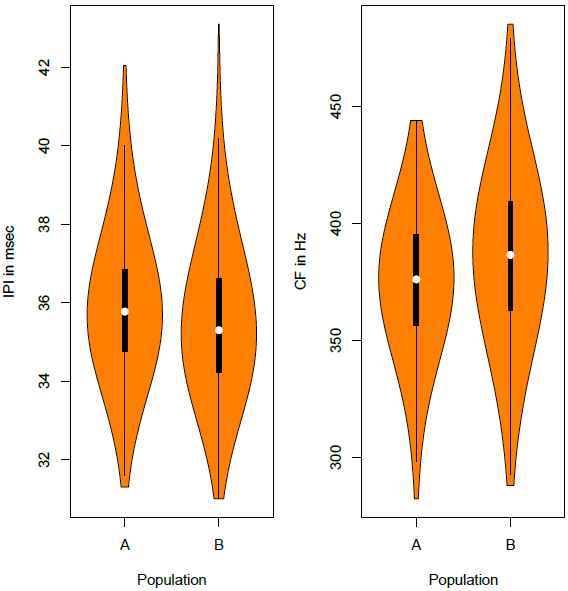
RIL phenotypes. Violin plots showing the distribution of average trait values for 840 lines from the A population and 816 lines from the B population. White circles show the median, outer quartiles are indicated by the thick line, and range excluding outliers is shown by the thin line; envelope width shows the density curve. Samples sizes per line range from 4 to 71 (average = 16).

**Figure S2.**
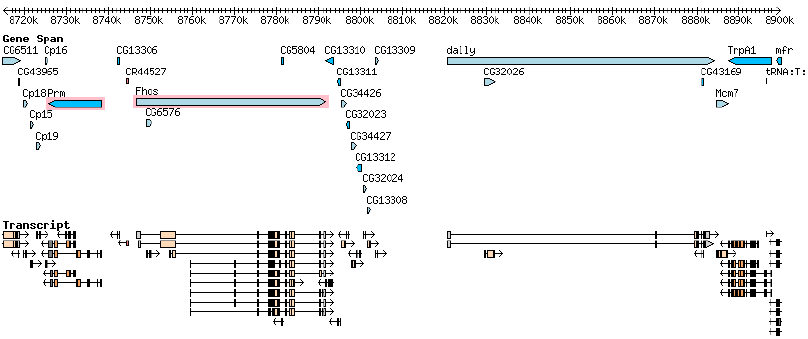
Genes at a 3L QTL. This figure shows all of the genes annotated within the 2-LOD drop confidence interval of our most significant QTL: the CF QTL in population A peaking at 8.78 Mb (QTL 1 in Table 1). Gene spans are shown in light or dark blue, depending on orientation, with gene models shown below in orange. The two genes we consider the best candidates (*Prm* and *Fhos*) are outlined in pink. Image is from the UCSC Genome Browser.

**Figure S3.**
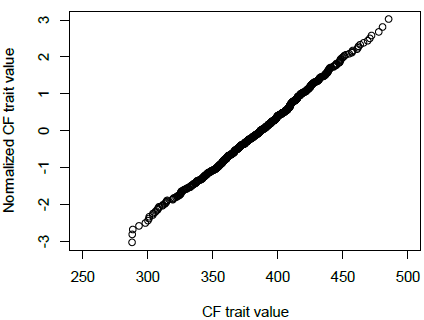
Normalization of carrier frequency. This figure illustrates the effect of t-rank normalization on the CF trait. Each point is the median value for one recombinant strain. Note that the main effect of normalization is on outliers. Only the B population is shown, but A is very similar.

**Figure S4.**
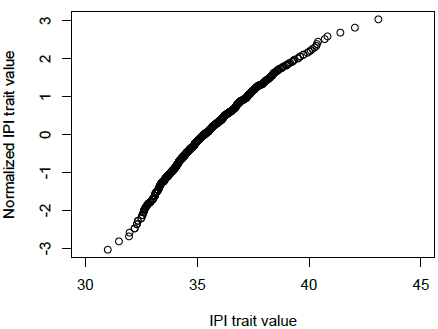
Normalization of inter-pulse interval. This figure illustrates the effect of t-rank normalization on the IPI trait. Each point is the median value for one recombinant strain. Note that the main effect of normalization is on outliers. Only the B population is shown, but A is very similar.

